# Accelerated growth increases the somatic epimutation rate in trees

**DOI:** 10.1101/2024.05.07.592680

**Authors:** M Zhou, G Schmied, M Bradatsch, G Resente, R Hazarika, I Kakoulidou, M Costa, M Serra, E Uhl, RJ Schmitz, T Hilmers, A Toraño Caicoya, A Crivellaro, H Pretzsch, F Johannes

**Affiliations:** Plant Epigenomics, Technical University of Munich, Freising, Germany; Chair for Forest Growth and Yield Science, Technical University of Munich, Freising, Germany; Department of Agricultural, Forest and Food Sciences, University of Torino, Grugliasco (TO), Italy; Department of Botany and Landscape Ecology, University of Greifswald, Greifswald, Germany; Department of Genetics, University of Georgia, Athens, GA, USA

**Keywords:** Fagus sylvatica, DNA methylation, epimutation rate, somatic epigenetic drift, somatic evolution, somatic epimutation, tree epigenomics

## Abstract

Trees are critical components of ecosystems and of major economic importance. Due to their extraordinary longevity and well-defined modular architecture they have also emerged as model systems to study the long-term accumulation of somatic mutations in plants. Coupled with retrospective life-history and environmental data, trees can offer unique insights into mutational processes that would be difficult to obtain with prospective studies. In addition to genetic mutations, somatic epimutations in the form of stochastic gains and losses of DNA cytosine methylation have been shown to accumulate rapidly during ontogeny. One hypothesis is that somatic epimutations originate from DNA methylation maintenance errors during mitotic cell divisions, which would predict that their rate of accumulation scales with growth rate, rather than with age.

Here we test this hypothesis in European beech. We analyzed one of the oldest continuously measured experimental plots in the world. The plot contains an even-aged beech stand that was established in 1822 and monitored for growth at regular intervals until the present. Starting ∼150 years ago, alternative thinning strategies were applied to subplots of this experiment, resulting in differential stem growth rates among trees. We show that accelerated growth significantly increased the per-year somatic epimutation rate at CG dinucleotides, and that this effect is accompanied by differences in cell division rates. Hence, faster growth elevates the chances for methylation maintenance errors during DNA replication per unit time. As somatic CG epimutations can be stably inherited to subsequent generations in plants, our insights have direct ecological and evolutionary implications.

---

Trees are among the longest-living organisms on earth. They have critical ecosystem functions, and continue to be of major economic importance (1). Perhaps due to their longevity, sessile life-style and modular nature, many tree species have evolved a remarkable degree of phenotypic plasticity in response to environmental stressors (2). Molecular evidence suggests that these plastic responses are at least in part driven by epigenetic mechanisms (3). One example is priming in Norway spruce (*Picea abies*), where hormonal exposure of seedlings can induce an epigenetically-encoded stress memory that facilitates more effective resistance to insect attacks later in life (4). Although such epigenetic memories are typically lost with passage into the next generation (3), their ecological consequences are highly relevant, given the ontogenetic time-scales involved (5). In addition to transient epigenetic effects that function as part of a stress response, there are also stable epigenetic changes that occur stochastically during development as a result of epigenome maintenance failures (6). A well-characterized form of such changes are accidental gains and losses of DNA cytosine methylation, a phenomenon that has been termed ‘spontaneous epimutation’. Numerous reports in plants indicate that epimutations occurring at CG dinucleotides are faithfully inherited mitotically and even meiotically (7–19).

When CG epimutations originate in the shoot apical meristem (SAM), a small population of stem cells at the shoot apex, they often become fixed in lateral branches via somatic epigenetic drift (20). This follows from the cell biology underlying lateral shoot formation: cell lineages that initiate lateral shoots are derived from a small number of precursor cells at the SAM periphery (21). The repeated “sampling” of precursors creates a series of strong cellular bottlenecks with each branching event (20, 22). As a result, somatically fixed epimutations tend to appear at high frequencies in lateral organs such as fruits or leaves, and are therefore detectable using sequencing approaches (**see note in *Methods***). Recent estimates in *Populus trichocarpa* show that fixed somatic epimutations accumulate at a rate of 8.0×10^−6^ per CG per haploid genome per year (12). This rate is about four orders of magnitude higher than the somatic genetic mutation rates found in *P. trichocarpa* (2.66×10^−10^) (12), *Quercus robur* (2.5×10^−10^), *Shorea laevis* (3.8×10^−10^) and *Shorea leprosula* (4×10^−10^).

There has been substantial interest in trying to understand the molecular mechanisms underlying the formation of somatic CG epimutations, as well as the factors that determine their rate of accumulation (6). Ultimately, these rates must reflect the accuracy of the maintenance methyltransferases that act on CG dinucleotides. In a simple molecular model, the stochastic loss of CG methylation occurs due to the imperfect enzymatic activity of METHYLTRANSFERASE 1 (MET1) (23), which is the primary methyltransferase that targets CG sites. During DNA replication, hemimethylated CGs are detected by the VARIANT IN METHYLATION protein family, which recruits MET1 to deposit CG methylation on the newly synthesized strand by way of “template copying” (24). Accidental failures of MET1 to carry out this catalytic step can lead to permanent methylation losses in daughter cells and their descendant lineages. Homologous to mammalian systems (25, 26), MET1 also appears to have a *de novo* function in plants (12, 18), which seems to be a source of stochastic gains of CG methylation. In addition to replication-dependent maintenance, a subset of CG dinucleotides is also redundantly targeted for methylation by the so-called RNA-directed DNA methylation pathway, which acts in the non-replicative part of the cell cycle (27).

Under a replication-dependent maintenance model, the rate of accumulation of somatic CG epimutations should scale with growth rate (i.e. growth per unit time). This has become known as the “cell-division hypothesis” (28). An alternative model is that the rate of accumulation scales with age, rather than with growth rate (29). The “age-related hypothesis” implies that CG methylation errors arise mainly in a replication independent manner. Interestingly, studying somatic genetic mutations in two tropical tree species, *S. laevis* (slow-growing) and *S. leprosula* (fast-growing), Satake et al. (29) recently found strong support for the age-hypothesis. Their analysis showed that the rate of somatic genetic mutations per meter of growth was 3.7-fold higher in *S. laevis* than in *S. leprosula*. This difference in the mutation rate appeared to scale with the slower stem growth rate of *S. laevis* compared to *S. leprosula*, thus yielding a constant somatic mutation rate per year between the two species. Whether these conclusions hold for the accumulation of CG epimutations remains unknown.

Here we provide a direct empirical test of this question. Using European beech (*Fagus sylvatica* L.) as a model, we took advantage of one of the oldest continuously measured experimental plots in the world. The plot contains an even-aged beech stand that was established in 1822 and monitored for growth at regular intervals until the present. Starting about 150 years ago, different thinning strategies were applied to subplots of this experiment, which has resulted in differences in the average stem growth rates of trees among subplots. Epimutation comparisons between slow and fast growing trees revealed strong support for the cell-division hypothesis. We found that a 1.79 fold increase in stem growth rate (cm^2^ yr^−1^) was associated with a 1.66 fold increase in the somatic CG epimutation rate within gene body methylated (gbM) genes, which suggests that this effect is likely mediated by differences in cell division rates.

Our results provide the first demonstration that accelerated growth contributes to the formation of somatic CG epimutations. As somatic CG epimutations are often inherited across generations in plants, our results highlight an interesting example of how developmental processes occurring at the level of individual trees can shape epigenetic diversity at the level of populations in the long-term.

## Results

### Experimental Stand Thinning Leads to Accelerated Growth

The Fabrikschleichach 15 (FAB15) long-term thinning experiment in European beech is one of the oldest continuously monitored experimental plots (***Methods***). The ∼0.4 ha sized unthinned, moderately thinned, and heavily thinned plots of FAB 15 were established in 1870/1871 in a 48-year-old, even-aged European beech stand that originated from natural regeneration by shelterwood cutting of a beech forest in 1822 (30). The managed plots were thinned 12 times since their establishment with a significant impact on stand growth dynamics (31). In 2020, 138 trees remained in the unthinned, 70 in the moderately thinned, and 40 in the heavily thinned plot. This resulted in different average stem growth rates of 8.1 cm^2^ yr^−1^ (unthinned), 12.2 cm^2^ yr^−1^ (moderate), and 16.4 cm^2^ yr^−1^ (heavy) within the period from 1822 to 2020, which differed significantly from each other (*P* ≤ 0.0001 for all three comparisons with pairwise Wilcoxon rank sum test and Bonferroni correction). The average stem growth rate of our sample trees (see below) were 12.2 cm^2^ yr^−1^ (tree 109, moderately thinned) and 21.8 cm^2^ yr^−1^ (tree 171, heavily thinned). Hence, thinning of beech stands promotes accelerated growth of single trees.

### Accelerated Growth Is Accompanied by Increased Xylem Cell Counts

We selected two focal trees from the moderately (tree 109) and heavily thinned (tree 171) plots to further study the impact of differential thinning on growth characteristics. These two trees were representative for the different thinning intensities in terms of growth dynamics (**Fig. 1 *A* and *B***). By the time of the last measurement in 2020, tree 171 was only slightly taller than tree 109 (42.8 m vs. 42.4 m), but was ∼1.3 times thicker at diameter breast height (DBH: 72.3 cm vs. 55.1 cm), although both trees had been planted in the same year (**Fig. 1*C***). The differential stem growth rates are reflected by the growth trajectories of both trees (**Fig. 1*D***). To evaluate whether differences in stem growth rate are the result of cell size expansion or increased cell proliferation, we performed cell count assays of xylem vessels, fiber cells, and parenchyma cells. On average, tree 171 had significantly more cells per annual ring than tree 109 (*P* ≤ 0.0001, **Fig. 1 *E*-*G***), suggesting that growth acceleration is accompanied by an increased cell division rate per unit time. Although these cell count data provide no direct information about cell division rates of other cell types, most notably the stem-cell compartment of the shoot apical meristem (SAM), they do show that cell proliferation is, in principle, affected by accelerated growth.

**Fig. 1.**
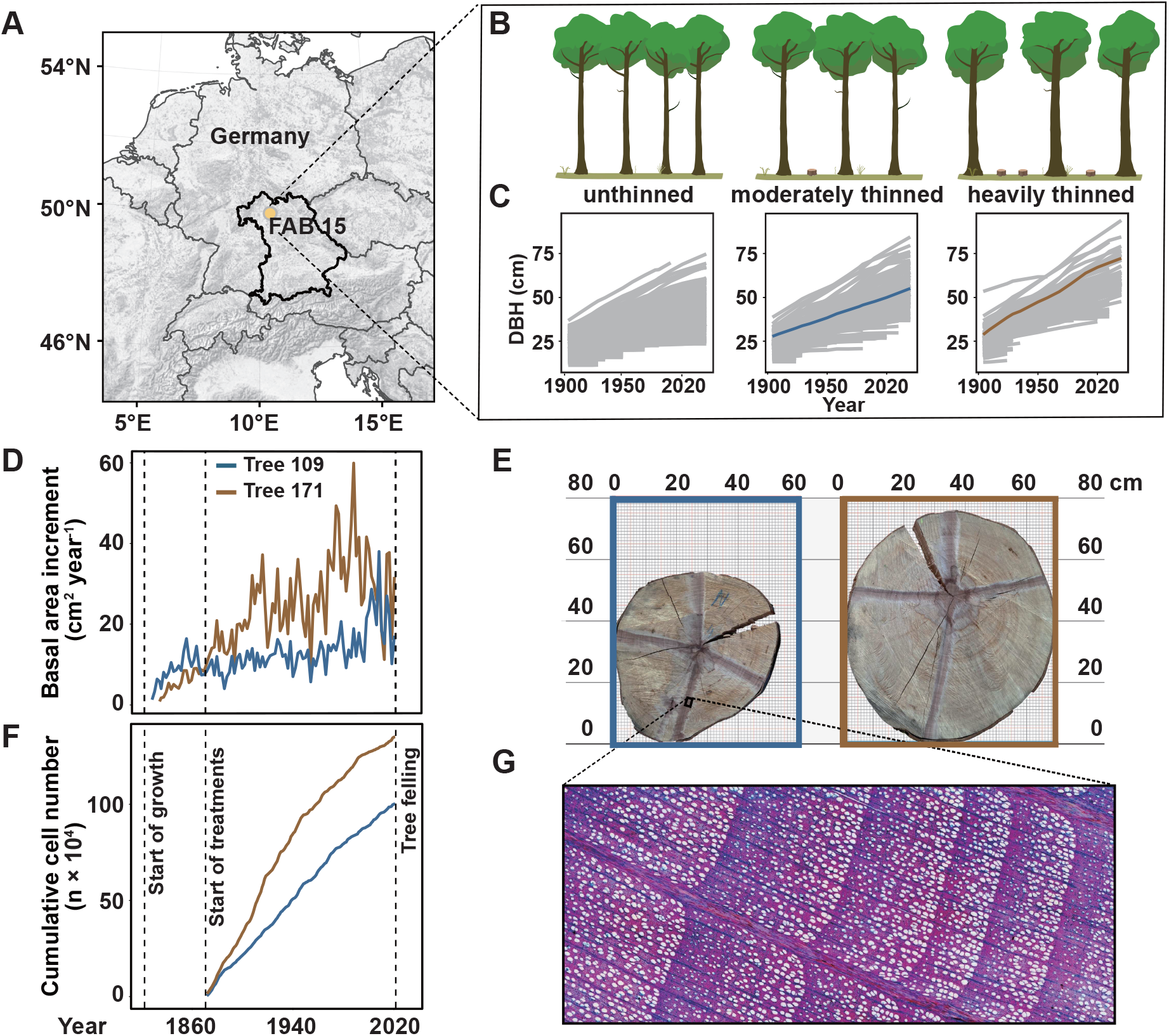
Overview of the location (**A**) and treatment design (**B**) of the FAB15 long-term thinning experiment in European beech. The different thinning intensities resulted in different tree dimensions (**C**,**E**) and growth trajectories (**C**). We relied on representative sample trees from the moderately (tree 109) and heavily thinned (tree 171) plots to estimate differences in somatic epimutation rates, growth rates (**D**) and cumulative cell production (**F**), the latter derived from wood anatomical analyses (**G**).

### Accelerated Growth Increases the Genome-Wide Somatic Epimutation Rate

The cell-division hypothesis predicts that accelerated growth increases the somatic epimutation rate per unit time. By contrast, the age-related hypothesis predicts no such effect. To test this, we generated whole genome bisulfite sequencing (WGBS) data from leaves of tree 171 (N=10) and 109 (N=7). The leaf samples were chosen to broadly survey the 3D branching topology of each tree (**Fig. 2*A***). To be able to detect spontaneous epimutations that emerge at individual CG dinucleotides as well as in larger (∼100-200 bp) genomic regions, we identified single methylation polymorphisms (SMPs) and differentially methylated regions (DMRs) between samples using jDMR (11) (***Methods***). Clustering the samples based on their SMP or DMR profiles recapitulated the known branching topology of each tree (**Fig. 2*A* and Fig. S1, Table S2**), indicating that our SMP/DMR calling approach was successful at detecting somatically fixed CG epimutations. In addition to DNA methylation profiling, we also determined the ages of the sampled branches from branch disks (***Methods***). Together, these data allowed us to calculate DNA methylation divergence as a function of divergence time (in years) between all leaf pairs (**Fig. 2 *B* and *C***). Consistent with recent work in *P. trichocarpa (12, 16)*, CG methylation divergence increased gradually with divergence time in both trees. This was true at the level of individual CGs as well as at the level of regions (**Fig. 3*D***). However, divergence was visibly more rapid in tree 171 than in 109 (**Fig. 3*D***), suggesting that tree 171 accumulated somatic epimutations at a faster rate.

**Fig. 2.**
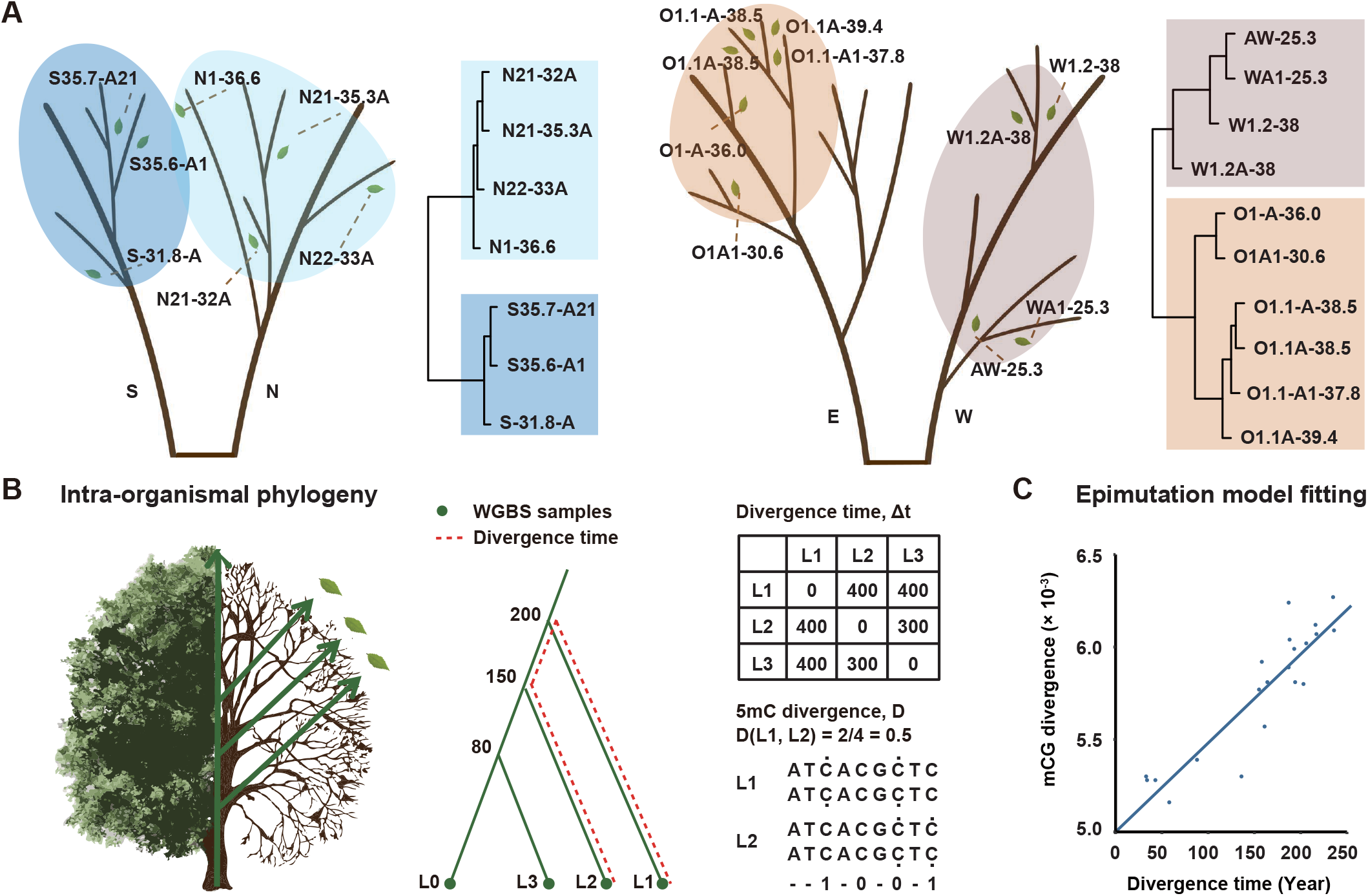
DNA methylation-based clustering of leaf samples recapitulates known tree topology. (**A**) Sample locations of leaves within the branching topologies of trees 109 (left) and 171 (right). Unsupervised sample clustering based on detected SMPs recapitulates the branching topologies of each of the two trees, indicating that SMP analysis identified somatically fixed epimutations. (**B**) A schematic showing that a tree can be interpreted as an intra-organismal phylogeny. The topology is given and the branch points and branch lengths can be dated by coring. For simplicity, only three branches are highlighted (L1, to L3) as well as the main stem (L0). Leaf WGBS measurements can be obtained and used to calculate pairwise DNA methylation divergence. Similarly, divergence times (in years) for pairs of leaves can be calculated by tracing back the ages of the branches to their most recent common branch point; here shown for L1 and L2. This can only be done down to the tree’s earliest branch point (in this case, Yr = 200) but not to earlier time points. (**C**) DNA methylation divergence increases as a function of divergence time. Divergence increases according to a neutral epimutation process, and depends only on the stochastic methylation gain and loss rates at the level of individual CG sites or at the level of regions.

**Fig. 3.**
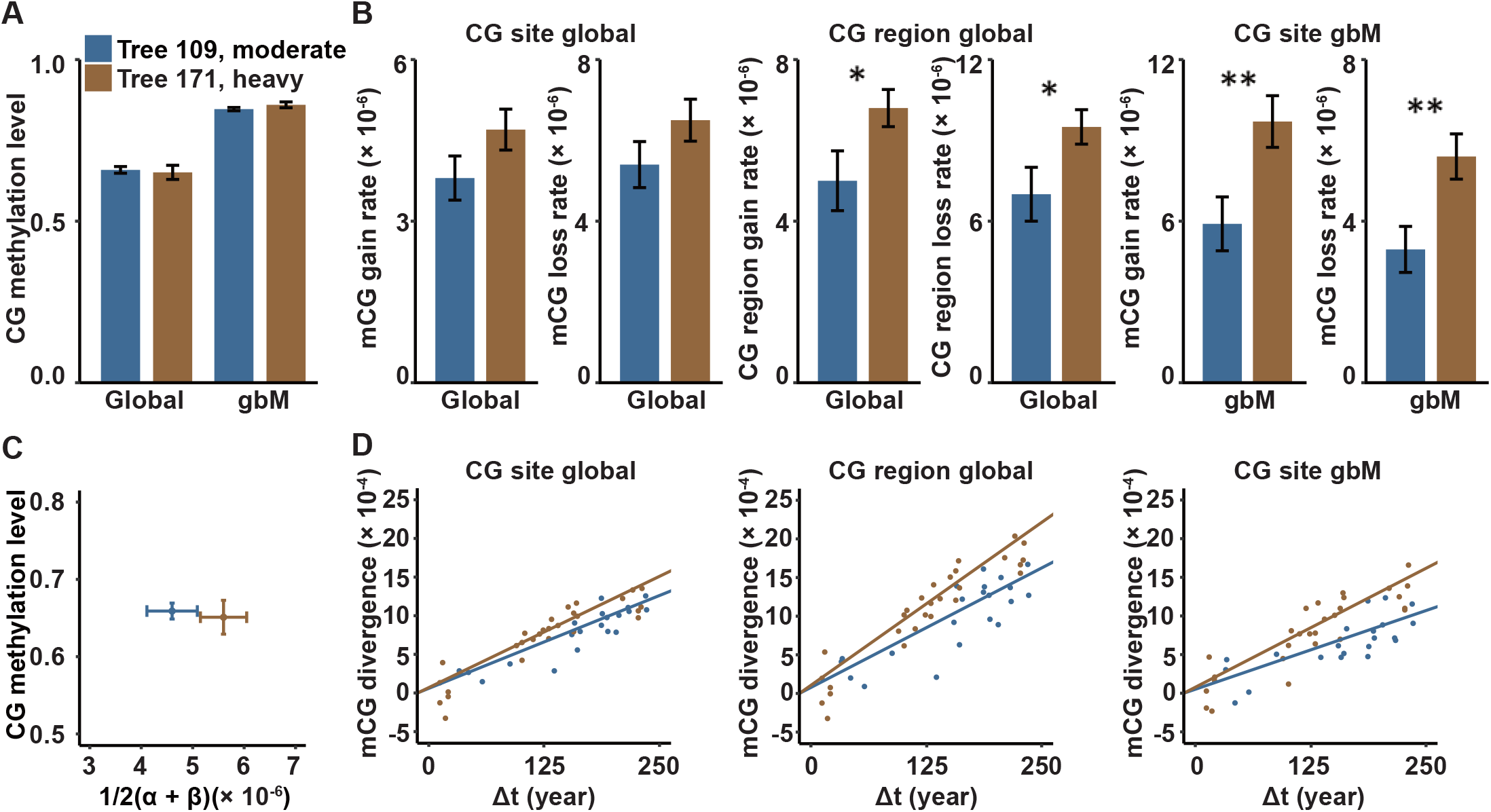
Analysis of somatic epimutation rates. (**A**) Comparison of CG methylation levels of gene body methylated (gbM) genes as well as genome-wide (global) reveal no significant differences between tree 171 (heavily thinned) and tree 109 (moderately thinned). However, tree 171 has a significantly higher per-site and per-region (**B**) CG epimutation rate than tree 109, both genome-wide (globally) as well as within gbM genes. *P*-values were calculated from Student’s *t* tests. Hence, very similar steady-state CG methylation levels can be accompanied by very different CG epimutation rates (**C**). (**D**) The epimutation rates differences between tree 171 and 109 are reflected in the fast CG methylation divergence as function of divergence time (in years). \**P* < 0.05; \*\**P* < 0.01.

To obtain direct estimates of these rates, we employed AlphaBeta (16). The software treats the tree branching topology as an intra-organism phylogeny of somatic cell lineages and fits an epimutation model to the mCG divergence data. At the genome-wide scale, the spontaneous mCG gain rate (α) and loss rate (β) were ∼1.22 times higher in tree 171 than in 109 on average (tree 171: a = 4.69×10^−6^ (SE = 0.38×10^−6^), β = 6.50×10^−6^ (SE = 0.52×10^−6^) per CG per haploid genome per year; tree 109: a = 3.83×10^−6^ (SE = 0.41×10^−6^), β = 5.36×10^−6^ (SE = 0.57×10^−6^) per site per haploid genome per year (**Fig. 3*B* and Table S3**). A similar picture emerged for region-level epimutation rate estimates (tree 171: a = 6.79×10^−6^ (SE = 0.46×10^−6^), β = 9.45×10^−6^ (SE = 0.64×10^−6^) per 100 bp region per haploid genome per year; tree 109: a = 4.99×10^−6^ (SE = 0.74×10^−6^), β = 6.96×10^−6^ (SE = 1.03×10^−6^) per 100 bp region per haploid genome per year (**Fig. 3*B* and Table S3**). Interestingly, despite these epimutation rate differences, genome-wide steady-state CG methylation levels were very similar between the two trees (**Fig. 3 *A* and *C***). This observation can be attributed to the nearly proportional increase in gain and loss rates in tree 171 relative to 109, which theory predicts to result in unchanged steady-state methylation (9, 16, 32)(***Methods***).

### Accelerated Growth Increases the Somatic CG Epimutation Rate in GbM Genes

The above genome-wide epimutation rate analysis could include subsets of CG dinucleotides that are redundantly targeted by *de novo* methylation pathways (27). The methylation gain and loss dynamics underlying our rate estimates may therefore be independent of the number of cell divisions. We sought to test if our conclusions still hold when focusing on CG sites that are exclusively targeted by MET1, whose maintenance activity is restricted to DNA replication. CG sites in gene body methylated (gbM) genes provide an ideal framework to assess this. gbM genes are evolutionary conserved genes in plants that feature high CG methylation and virtually no non-CG methylation (i.e. methylation in context CHG and CHH (where H = A, T, C)) (33–36). GbM genes are enriched in housekeeping functions, are moderately expressed, and display low transcriptional variability across cells and tissues (37–40) (extensively reviewed in Refs. (41, 42). Recent work with *A. thaliana* DNA methylation mutants revealed that the CG epimutation rate in gbM genes is not strongly affected by loss of function mutations in DNA methylation pathways other than MET1 (11, 18). We identified gbM genes in each leaf sample (***Methods***). This resulted in a liberal set of ∼10,000 gbM genes out of the 65,000 annotated genes and pseudogenes in the current European Beech (*Fagus sylvatica* L.) reference assembly. Using CGs extracted from gbM genes, we repeated our epimutation rate estimation. The analysis revealed that the rate difference between trees 171 and 109 became even more pronounced, with CG epimutation rates being ∼1.66 times higher in tree 171 than in tree 109 on average (tree 171: a = 9.66×10^−6^ (SE = 0.96×10^−6^), β = 5.62×10^−6^ (SE = 0.56×10^−6^) per CG per haploid genome per year; tree 109: a =5.94×10^−6^ (SE = 1.02×10^−6^), β = 3.32×10^−6^ (SE = 0.57×10^−6^) per site per haploid genome per year) (**Fig. 3*B* and Table S3**). These results provide further, albeit indirect, evidence that the epimutation rate differences are likely coupled with an increased cell division rate of the SAM and/or the cell lineages leading up to the formation of leaf primordia.

## Discussion

Knowledge of the factors that modulate the rates of somatic (epi)mutations in trees are of fundamental interest to plant biology. Somatic epimutations that arise in the form of stochastic gains and losses of cytosine methylation have been shown to accumulate several orders of magnitude faster than genetic mutations within the tree branching architecture. Molecular models suggest that these stochastic events are the results of DNA methylation maintenance mistakes during cell divisions. According to this model, an increase in growth rate should lead to faster accumulation of somatic epimutations per unit time.

Using European beech as a study system, we have provided a direct experimental test of this hypothesis. By estimating epimutation rates in fast and slow growing, same-age beech trees, we found that a 1.79 fold increase in stem growth rate (cm^2^ yr^−1^) led to a 1.66 fold increase in the somatic CG epimutation rate within gene body methylated (gbM) genes. Since our analysis focused on fixed CG epimutations in leaves, these rate differences likely stem from differences in the number of cell divisions of the SAM and / or the cell lineages leading up to the formation of leaf primordia. This is corroborated by our focus on CG sites that are exclusively targeted for methylation maintenance by methyltransferase MET1, which is known to act in a DNA replication-dependent manner. Our results thus provide strong evidence that fast growing trees experience more cell divisions per unit time, which in turn increases the chances for DNA methylation maintenance errors. A direct assessment of the relationship between stem growth rate and the rate of cell divisions of the SAM is experimentally challenging, particularly in *vivo* (43). In *vitro* cell proliferation assays of the SAM could be performed (44), which may or may not reflect the in vivo situation.

Our results are at odds with a recent report of the somatic mutation rates in two tropical tree species, which was shown to be mediated entirely by age, rather than growth rate. This result would suggest that all fixed somatic mutations arise from errors in replication-independent DNA repair, rather than during DNA synthesis. While there is evidence from the mammalian field that age does have an effect on somatic mutation rates, for example in oocytes (28), the lack of a contribution from growth rate should be further examined. In contrast to our experimental design, which used even-aged trees, the recent study in tropical trees varied both age and stem growth rate at the same time. Ruling out an effect of growth rate would therefore require a direct demonstration that the cell division rates of the two species are indeed the same. Although this was demonstrated for fiber cells in these species (29), the dynamics of these cell types may not be directly related to the cell lineages that form the leaves from which the rate of fixed somatic mutations had been inferred.

In our study, the “cleanest” subset of CG sites that could provide clues about methylation errors stemming for DNA replication are CGs within gene body methylated genes. The fact that the epimutation rates at these sites were modulated by stem growth rate is somewhat surprising, given our recent finding that the per-generation epimutation rate in these regions is extremely robust to environmental stress and genetic variation (17), both of which are known to affect growth rate parameters. We hypothesize that – while the per unit time rate can vary – the per-generation epimutation rate is tightly controlled. This would imply that accelerated growth in trees is offset by early growth cessation, senescence or mortality (45, 46). We recently uncovered a similar trade-off when comparing the annual plant *A. thaliana* with the *P. trichocarpa* (16). Although the per unit time (i.e. per year) CG epimutation rate in *P. trichocarpa* was about 2 orders of magnitude lower than *A. thaliana*, the per-generation epimutation rate was remarkably similar (put rates 10^−4^). This effect may be related to the hypothesis that long-lived perennials have evolved mechanisms to limit the number SAM cell divisions (per unit time) as a way to slow “mutational meltdown” (47). However, these arguments do not hold for species that lack well-defined generations. These include species that propagate clonally through the formation of stolons, tillers or rametes. Questions arise as to how differential growth rates among clones, if they exist, relate to patterns of natural epigenetic diversity as well as to population-genetic inferences that are based on these patterns. Our study provides the needed rationale to begin to explore such questions.

## Methods

### Study Site and Sampling Design

We relied on the long-term thinning trial Fabrikschleichach 15 (FAB15) in a European beech (*Fagus sylvatica* L.) forest in Central Europe (**Fig. 1*A***) to obtain trees with exactly the same age, but with strongly divergent growth rates. The region is characterized by a mild climate with an average annual temperature of 7.5 °C and an annual precipitation of 820 mm, with the natural vegetation being submontane European beech-sessile oak forests. FAB15 is one of the longest continuously measured forest experimental sites worldwide and consists of three differently managed plots, each with a size of ∼0.4 ha: unthinned, moderately thinned, and heavily thinned. The beech forest, now over 200 years old, originated from a shelterwood cutting in 1822, and the various treatments and continuous measurements began in 1870/1871. The unthinned plots remained essentially untouched, apart from the occasional removal of dying or dead trees to prevent possible damage to the stand from fungal or insect infestation. In contrast, the managed plots were moderately or heavily thinned to reduce stand density by removing mainly suppressed trees, but also tall trees, especially in the case of the more intense treatment (31). In total, the managed plots have been thinned 12 times with different intensities since their establishment, resulting in different tree sizes and stem growth rates (**Fig. 1 *B* and *C***). We selected a representative, average sample tree from each thinned plot for further analyses, which was felled and precisely measured. Both selected trees had a particularly low branching point to allow for distinct differences in branch divergence time within the same tree. After felling, branches along the crown periphery were selected (distributed across lower, middle, and upper crown) and leaves were sampled from the ends of these branches (**Fig. 2*A***), which were kept frozen at -80 °C for subsequent DNA methylation analysis. For a correct assignment of branches within the tree architecture, we reconstructed the tree topology and only selected those branches for sampling whose branching path could be clearly traced back. Furthermore, we obtained stem and branch disks from different positions along the stem axis and crown topology to allow for age and stem growth rate estimations. In more detail, a stem disk was extracted from each tree at 1.3 m height, while branch disks from leaf-sampled branches were taken 10 cm before and after each branching position.

### Growth Rate Analysis

We used the stems disks to measure the annual tree ring widths from four cardinal directions (N, E, S, W; see **Fig. 1*E***) to the nearest of 1/100 mm using a digital positioning table for stem disk measurements (Digitalpositometer, Biritz and Hatzl GmbH, Austria) after sanding the disks with progressively finer sandpaper (up to 800 grit). Tree ring widths were transformed to basal area increments (bai) using the formula 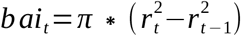, where *r*represents the tree’s radius at 1.3 m height and the respective year *t*. We defined the stem growth rate as the average annual basal area increment, considering all four series per stem disk.

### Branch Age Determination

For determining the age of the different branches, we utilized the derived branch disks, which were sanded to enhance the visibility of annual tree ring borders (see also stem disk preparation). Tree rings of all branch disks (from four cardinal directions per disk) were measured with the digital positioning table Lintab 5 and the software TSAPWin (both Rinntech, Heidelberg, Germany). We visually cross-dated all tree ring series, taking into account distinctive growth patterns across the different samples to ensure the correct dating of the individual tree rings (48). We counted all tree rings from bark to pith as an estimation of the age of each branch.

### Cell Count Assays

We relied on wood anatomical methods to derive estimates of annually produced xylem cells for both sample trees. Stem disks were cut with a small circular saw into ∼1cm wide sections, capturing all tree rings from the pith to the bark. For both trees, we used a section without any cracks or damages for further wood anatomical analyses. The cross sections were cut into smaller segments of 3-5 cm before taking transverse micro sections of 10-20 µm thickness with a sliding GSL-1 microtome (Schenkung Dapples, Zürich, Switzerland). Sample preparation followed a protocol by Gärtner & Schweingruber (49). In more detail, all samples were bleached and washed with distilled water, before they were double-stained with safranin and astrablue for 5 minutes (mixture of 1:1). Subsequently, the samples were again rinsed with distilled water, ethanol of increasing purity (80 %, 96 %, and anhydrous ethanol) for dehydration, and xylene before being permanently embedded on a glass slide with Canada balsam and dried out in an oven at 60°C for 24h. We used an optical microscope (Zeiss Axio Imager Z2) at × 200 magnification, equipped with an integrated camera (Zeiss Axiocam 305 color) to capture micrographs of wood anatomical features. The micrographs were stitched together using the software ZEN 3.2 blue edition (all three by Carl Zeiss Microscopy Deutschland GmbH, Oberkochen, Germany). Finally, we utilized CARROT, a software based on Deep Convolutional Neural Network (DCNN) algorithms, specifically programmed and trained for quantitative wood anatomy analyses. The software was employed to recognize and segment all types of cells for each ring from the obtained micrographs (50). We assessed the number of xylem cells per tree ring along a 5000 pixel wide band extending from the start of treatment in 1870 to the felling date in 2020.

### Whole Genome Bisulfite Sequencing Analysis

DNA was extracted from leaves using Qiagen’s DNeasy Plant Mini Kit. WGBS library preparation and 150bp paired-end Illumina sequencing was performed by the Beijing Genome Institute (BGI). The *Fagus sylvatica* L. reference genome and annotation was used from http://www.beechgenome.net/ (51). The WGBS data was processed with the MethylStar pipeline (52) for read trimming and alignments, and METHimpute for methylation state calling (53). The mean coverage of our samples was about 17X, and the mean mapping rate was about 70% (see **Table S1**).

### Identification of Gene Body Methylated Genes

We identified gene body methylated (gbM) genes using gbMine (github.com/jlab-code/gbMine). The software uses the outputs of MethylStar and an annotation (gff3) file. gbMINE provides two flags, -- genomicBackground and -- exons, which allows us to identify gbM genes using four different genomic features combinations. We only set -- exons, which means that we only consider exonic cytosines in both the foreground and the background. Using a binomial model, gbMINE classifies a gene as gbM if the proportion of mCG in the exons of that gene is statistically higher than that of exons in the background, while the proportion of mCHG and mCHH is not significantly different. We obtained a set of gbM genes for each sample, and used the union as a liberal set of gbM genes. As a further filter, we calculated the non-CG methylation levels of introns of the union set, which generated a distribution of intronic non-CG methylation levels for candidate gbM gene. Finally, we only retained gbM genes whose intronic non-CG methylation levels were less than the median of that distribution.

### SMP and DMR Calling

DMR calling was performed using jDMR (11) (github.com/jlab-code/jDMRgrid). Briefly, we divided the genome into windows of 100 bp each. Within these windows, we calculated the number methylated and total cytosines, which were used as input for Methimpute (53), a finite state Hidden Markov Model (HMM) with binomial emission densities. We employed Methimpute’s three-state HMM, where each 100 bp window was classified as methylated, unmethylated, or intermediate. The intermediate state calls are designed to capture somatic epiheterozygotes, which are only visible in WGBS data in the form of “intermediate” methylation levels. This approach yielded a matrix representing the methylation state of each region and sample (**Table S2**). jDMR queries this matrix to identify methylation state switches between samples within each 100 bp window (e.g. unmethylated to intermediate methylated) and defines these as DMRs. A similar strategy was employed to call SMPs, with the window size being reduced to single CG sites. Although our SMP and DMR analysis was based on bulk leaf methylome data, jDMR’s HMM approach is robust to measurement noise arising from the cell layer-specificity in which somatic variations often originate in the SAM (54, 55). This becomes evident from the clustering results in **Fig. 2*A***.

### Epimutation Rate Estimation

We estimated somatic epimutation rates using AlphaBeta (16). AlphaBeta requires methylation data and pedigree data as input. Following Shahryary et al. (16) and Hofmeister et al. (12), we treated the tree branching structure as a pedigree of somatic cell lineages, where the leaves represent the lineage end-points. AlphaBeta calculates the methylation divergence (*D*) and divergence time (Δ*t*) between all pairs of leaf samples. The model parameters are estimated using numerical nonlinear least squares optimization. We estimated epimutation rates under a neutral epimutation model (AlphaBeta’s ABneutralSOMA model).

## ACKNOWLEDGEMENTS

This work was partly funded by the Bundesministerium für Bildung und Forschung (Project: epiSOMA). We would like to thank the Bavarian State Institute of Forestry (Forest Protection Department) and the Ecophysiology group at TUM for their support with technical equipment and the Bavarian State Forests (BaySF) for their support in setting up and maintaining the underlying long-term experiments. Zhou M holds a fellowship from the China Scholarship Council (CSC NO. 202204910009).

