## Supplemental Figure 1 for "Accelerated growth increases the somatic epimutation rate in trees"

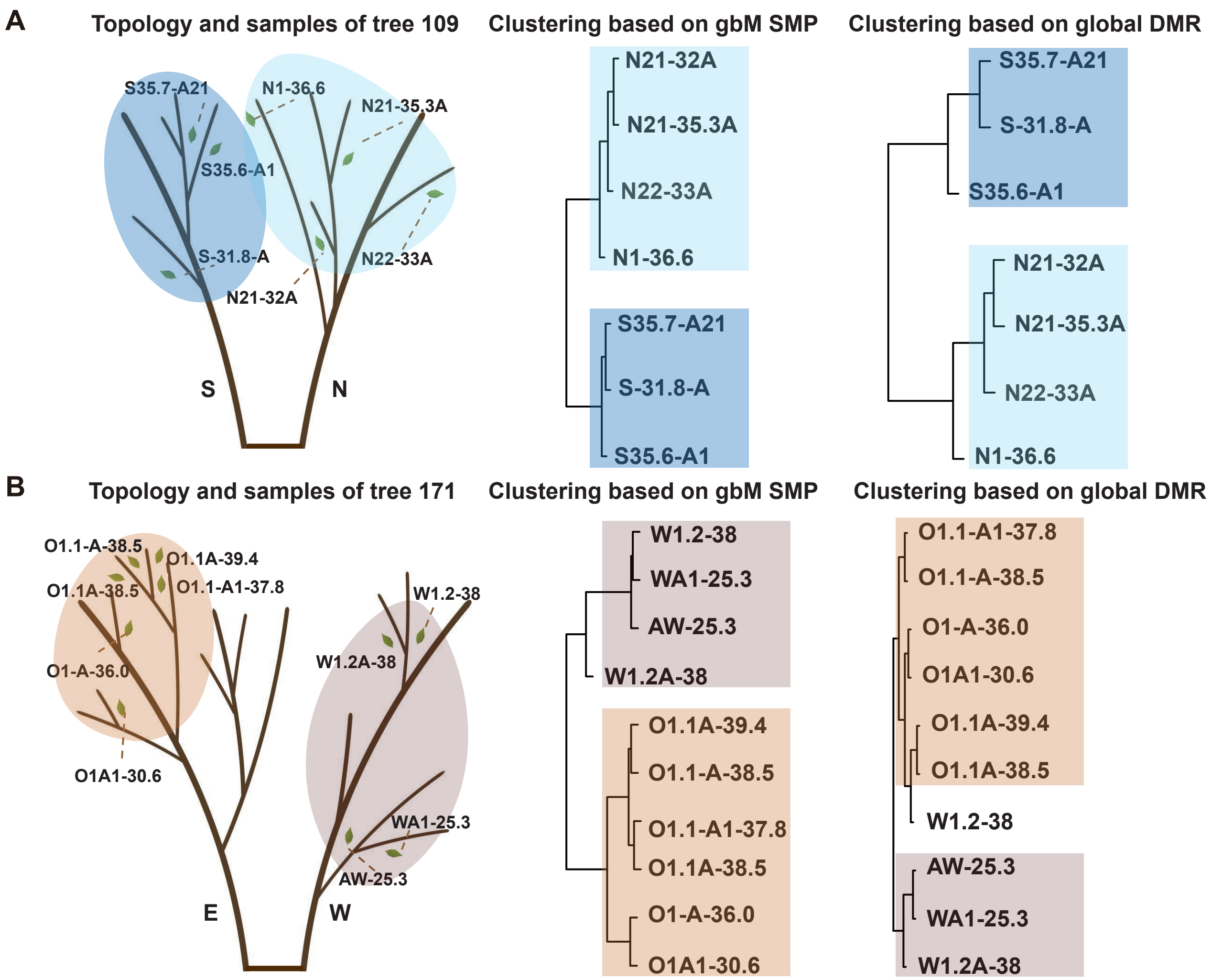


**Fig. S1.** DNA methylation-based clustering of leaf samples recapitulates known tree topology. (**A,B**) Sample locations of leaves within the branching topologies of tree 109 and tree 171. Unsupervised sample clustering based on detected SMPs on gbM genes and DMR recapitulates the branching topologies of each of the two trees.
